# Rapid receptor internalization potentiates CD7-targeted lipid nanoparticles for efficient mRNA delivery to T cells and *in vivo* CAR T-cell engineering

**DOI:** 10.64898/2026.01.23.701374

**Authors:** Jianhao Zeng, Tyler Ellis Papp, Awurama Akyianu, Alejandra Bahena, Lanfranco Leo, Faris Halilovic, Hamideh Parhiz

## Abstract

Targeted lipid nanoparticles (tLNPs) enable efficient mRNA delivery to T cells, allowing for *in situ* generation of chimeric antigen receptor (CAR) T cells without *ex vivo* manipulation. This strategy has shown promising therapeutical efficacy in preclinical studies of cardiac fibrosis, cancer, and autoimmune diseases. While multiple T-cell surface receptors have been targeted across studies for tLNP-mediated *in vivo* CAR T-cell generation and exhibit diverse efficiencies, their comparative performance and the mechanisms underlying these differences remain unclear. Here, we systematically compared tLNPs with antibody-based moieties targeting T-cell receptors including CD2, CD4, CD5, CD7, CD8, or a CD4/8 dual-targeting combination under identical conditions, assessing their mRNA delivery efficiency in human T cells and PBMCs *in vitro*, and subsequently validating the best performer *in vivo* in humanized mice. Among all moieties tested, CD7-targeting tLNPs achieved the highest mRNA delivery to T cells and efficiently generated functional CAR T cells *in vivo*. Mechanistic analysis revealed that receptor internalization, rather than the receptor abundance, is the primary determinant of delivery efficiency, a property intrinsic to each receptor and largely independent of antibody clone. These findings provide a rational framework for selecting optimal targeting moiety to enable highly efficient *in vivo* CAR T-cell engineering.

**Highlights:**

- Targeting CD7 outperforms other receptors for tLNP-mRNA delivery to T cells
- Receptor abundance does not predict tLNP-mRNA delivery efficiency
- Receptor internalization kinetics governs tLNP-mRNA delivery efficiency
- CD7-targeting LNP-mRNA enables efficient *in vivo* CAR T-cell engineering

**Graphical Abstract:** 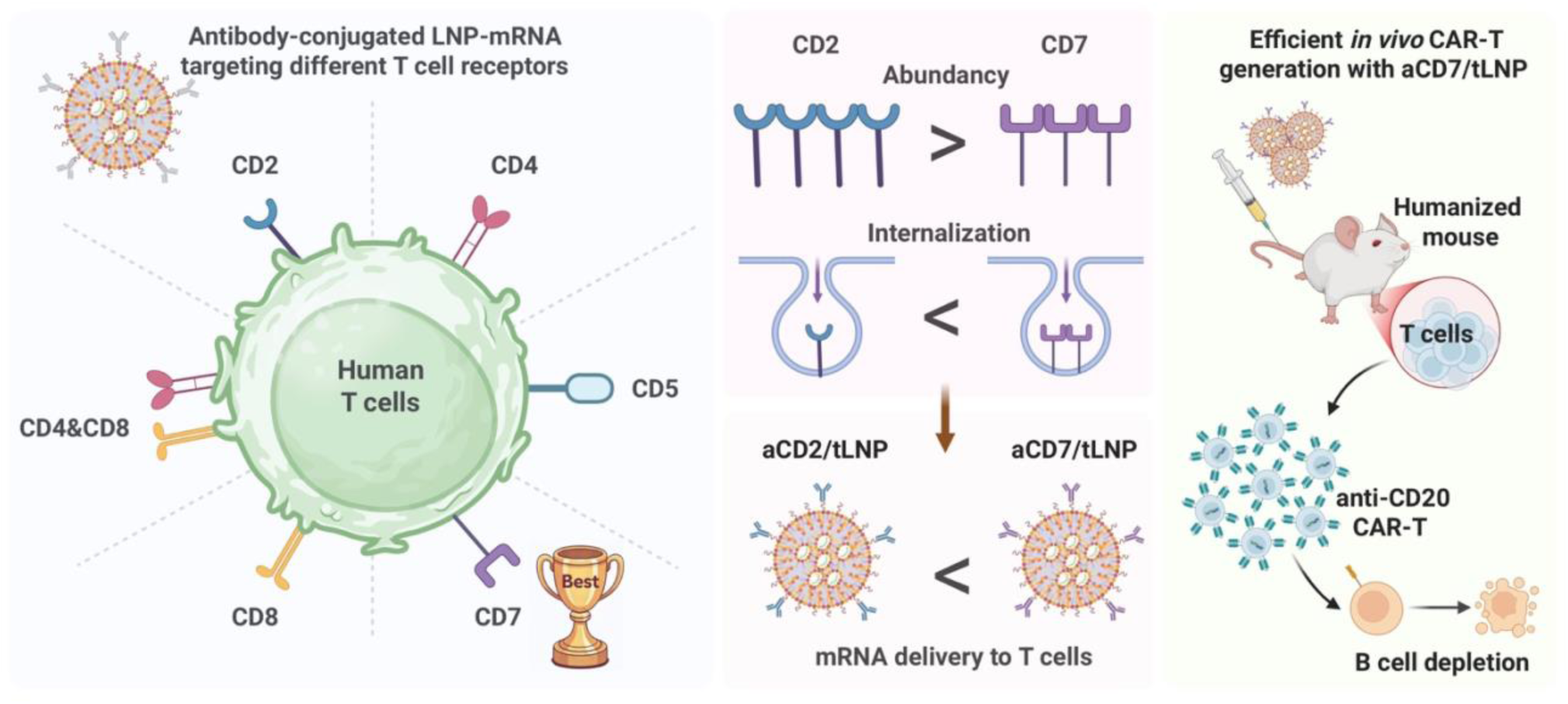

## Introduction

Messenger RNA (mRNA) delivered by lipid nanoparticles (LNPs) has emerged as a transformative platform for therapeutic development across vaccines, cancer treatments, and gene therapies[1–4]. A particularly compelling approach is the use of LNP-mRNA to generate chimeric antigen receptor (CAR) T cells directly *in vivo*. CAR T-cell therapy has revolutionized treatment of B-cell malignancies, delivering durable remissions and even cures [5–8]. However, current CAR T-cell manufacturing depends on complex *ex vivo* engineering workflows, requiring leukapheresis, viral transduction, expansion, quality control, and reinfusion, that are technically intensive, costly, and accessible only at specialized GMP facilities[9, 10]. Additional hurdles include the need for lymphodepletion conditioning via chemotherapy [11–13]. Prolonged CAR expression in virally induced T cells can drive sustained release of cytokines and long-term depletion of healthy B cells, leading to cytokine-release syndrome, extended immunosuppression, and increased infection [6, 14–16]. In contrast, *in vivo* generation of CAR T cells using LNP-mRNA can overcome many of these limitations. This approach eliminates the need for cell isolation and *in vitro* manipulation, thereby markedly simplifying the manufacturing process. Engineering endogenous T cells *in situ* also eliminates the requirement for chemo-conditioning and reduces associated toxicities. Moreover, mRNA enables the transient expression of CAR, mitigating risks associated with prolonged CAR T-cell activity and allowing for temporal control to minimize toxicity [17–19].

A major breakthrough enabling efficient *in vivo* mRNA delivery to T cells—and thereby enabling *in vivo* CAR T-cell generation—was the development of targeted lipid nanoparticles (tLNPs). Unlike macrophages or dendritic cells, T cells possess limited endocytic activity and do not efficiently uptake conventional unmodified LNP-mRNAs. The tLNP platform addresses this limitation by conjugating targeting moieties, such as monoclonal antibodies recognizing T-cell surface receptors, to the LNP surface to direct LNP-T cells engagement and trigger receptor-mediated endocytosis of LNP. This strategy enables potent and selective mRNA delivery to T cells following systemic administration [20–22]. tLNP-mediated *in vivo* CAR T-cell generation has demonstrated compelling efficacy across multiple preclinical and clinical studies. Rurik et al. demonstrated that aCD5/tLNPs enabled *in vivo* generation of anti-FAP CAR T cells, resulting in reversal of cardiac fibrosis in mice [22]. Hunter et al. reported that aCD8/tLNPs enabled *in vivo* generation of anti-CD19 CAR T cells and elicited an ‘immune reset’ in a nonhuman primate model of autoimmune disease [18]. Most recently, Wang et al. employed aCD8/tLNPs to generate anti-CD19 CAR T cells directly in patients with systemic lupus erythematosus, achieving robust B-cell depletion and reductions in pathogenic autoantibodies [23]. Across these studies, various T-cell surface receptors have been targeted by tLNPs, each yielding different levels of delivery efficiency [18, 21, 22]. However, a head-to-head performance comparison of tLNPs targeting these T cell receptors under identical experimental conditions has yet to be reported. Moreover, the mechanistic principles underlying the variable performance of tLNPs with different targeting moieties remain poorly defined. Addressing these questions would inform the selection of optimal targeting receptors and moieties for tLNP design to achieve robust mRNA delivery to T cells for in vivo CAR T-cell engineering.

Here, we systematically compared tLNPs targeting CD2, CD4, CD5, CD7, CD8, or a CD4/8 dual-targeting combination (referred to as CD4/8 hereafter) under uniform conditions. Using ZsGreen reporter mRNA as the tLNP payload, we quantitatively compared tLNP delivery efficiency *in vitro* in purified human T cells and in PBMCs, where T cells compete with other immune populaitons, followed by *in vivo* validation of the top-performing tLNP in humanized mice. To investigate the mechanistic principles underlying the tLNP delivery efficiency, we measured receptor abundance, assessed antibody-receptor internalization kinetics across multiple antibody clones, and quantified intracellular mRNA accumulation and resultant protein output. With the top performing tLNP, we further determined functional relevance by delivering anti-CD20 CAR mRNA to generate CAR T cells both *in vitro* and *in vivo*. We found that CD7-targeting tLNPs achieved the highest mRNA delivery to T cells and enabled potent CAR T-cell generation *in vitro* and *in vivo*. Mechanistically, the internalization capacity of the antibody–receptor complex, rather than receptor abundance, emerged as the primary determinant of mRNA delivery efficiency by tLNPs targeting that receptor. This internalization capacity appears to be an intrinsic property of each receptor and largely independent of the binding antibody clone. Together, our findings provide a framework for selecting optimal targeting moieties in designing tLNP-mRNA for efficient *in vivo* CAR T-cell engineering.

## Materials and Methods

### mRNA synthesis

mRNA was produced following previously described protocols [20, 24]. The amino acid sequences of encoded proteins (e.g., ZsGreen) were codon-optimized prior to generating the corresponding mRNA coding sequences. DNA templates for *in vitro* transcription, consisting of the T7 promoter, 5′ UTR, coding sequence, 3′ UTR, and poly(A) tail, were synthesized and cloned into the pUC-ccTEV-A101 plasmid vector. Plasmids were linearized and used as templates for *in vitro* transcription with the MEGAscript® T7 Transcription Kit (Invitrogen, Cat. AM133), according to the manufacturer’s instructions. Uridine triphosphate (UTP) was replaced with m¹ψ-5′-triphosphate (pseudouridine triphosphate; TriLink, Cat. N1081), and co-transcriptional capping was performed using CleanCap® (TriLink). Synthesized mRNA was purified using cellulose-based purification to remove double-stranded RNA contaminants, as previously described [25]. The integrity of purified mRNA was assessed by 2% agarose gel electrophoresis, and samples were stored at −20 °C until use.

### LNP-mRNAs preparation and characterization

mRNAs were encapsulated in lipid nanoparticles using the NanoAssemblr™ Ignite™ microfluidic formulation system (Cytiva) according to the manufacturer’s instructions. Lipids (SM-102, DSPC, cholesterol, and PEG2000) were dissolved in ethanol, and mRNA was diluted in 25 mM sodium acetate buffer (pH 5.0). The lipid solution and mRNA were combined within the microfluidic cartridge at a flow-rate ratio of 3:1. Following formulation, LNP–mRNA preparations were buffer-exchanged into 10% sucrose and concentrated to 1 mg/mL using a 10 kDa MWCO centrifugal filter (Millipore, Cat. UFC9010). Samples were stored at −80 °C until use. Encapsulated mRNA concentration was quantified using the Quant-iT™ RiboGreen assay (Thermo Fisher Scientific, Cat. R11490), as previously described [26]. Particle size, polydispersity index, and zeta potential of the LNP–mRNA formulations were measured by dynamic light scattering (DLS) using a Malvern Zetasizer instrument (Malvern, UK) following the manufacturer’s instructions.

### Antibody-conjugated LNP-mRNA preparation

To prepare targeted LNPs, monoclonal anti-CD2 (TS1/8, BioLegend, cat# 309202), anti-CD4 (A161A1, BioLegend, cat# 94222), anti-CD5 (UCHT2, BioLegend, cat# 98889), anti-CD7 (CD7-6B7, BioLegend, cat# 94122), anti-CD8 (SK1, BioLegend, cat# 92401), and anti-CD71 (B3/25, Bio X Cell, cat# BE036) were conjugated to LNP-mRNA using SATA-maleimide chemistry as described previously[27–29]. Briefly, LNP-mRNA was modified with maleimide functional groups (DSPE-PEG-mal) via post-insertion. The antibody was functionalized with SATA (N-succinimidyl S-acetylthioacetate) to introduce sulfhydryl groups for conjugation to maleimide. The sulfhydryl groups on the antibody were then conjugated to maleimide moieties through thioether chemistry. The resulting conjugates were purified using Sepharose CL-4B gel filtration columns (MilliporeSigma).

### Cell culture and medium

Human T cells and peripheral blood mononuclear cells (PBMCs) were obtained from STEMCELL Technologies and from the Human Immunology Core at the Perelman School of Medicine, University of Pennsylvania. Cells were cultured in RPMI 1640 supplemented with 10% fetal bovine serum (FBS), 100 U/mL penicillin, 100 µg/mL streptomycin, and 50 IU/mL recombinant human IL-2, and maintained at 37°C in a humidified incubator with 5% CO₂. T-cell activation was carried out using Dynabeads Human T-Activator CD3/CD28 (Thermo Fisher, cat# 11131D) following the manufacturer’s instructions.

### Flow cytometry analysis of tLNP-mRNA delivery efficiency

For cultured cells, one million cells were collected for flow cytometric analysis. For immune cells isolated from blood, peripheral whole blood was collected using heparinized micro-hematocrit capillary tubes, transferred immediately into blood collection tubes, and subjected to ACK red blood cell (RBC) lysis. For spleen samples, spleens were harvested, mechanically dissociated, passed through a 70-µm cell strainer, and treated with RBC lysis. One million cells per sample were used for flow staining. Cells were first stained with a live/dead viability dye (Invitrogen, cat# L34994) followed by Fc receptor blocking (BioLegend, cat# 422302). The T-cell staining panel included CD45-BUV805 (HI30, BD Biosciences, cat# 612891), CD3-APC (OKT3, BioLegend, cat# 317318), CD4-BUV395 (SK3, BD Biosciences, cat# 563550), CD8-BV421 (RPA-T8, BioLegend, cat# 301036), and the G4S linker antibody (E7O2V, Cell Signaling Technology, cat# 38907) for detection of CAR expression. For PBMC and spleen samples, additional antibodies included CD20-BV711 (2H7, BioLegend, cat# 302342), CD14-PE/Fire 810 (S18004B, BioLegend, cat# 399223), and CD56-BV605 (HCD56, BioLegend, cat# 318333). Sample acquisition was performed on an LSRFortessa flow cytometer (BD Biosciences), and data were analyzed using FlowJo software (v11)

### Antibody internalization assay

Activated human T cells were seeded into 96-well optical-bottom plates at 200,000 cells per well in 50 µL of T-cell medium. To promote cell adherence and minimize cell movement during imaging, plates were pretreated with 50 µL of 0.01% poly-L-ornithine per well for 2 hours at room temperature. Afterward, the solution was removed, and the plates were air-dried for 1-hour. Seeded cells were incubated for 2-hours before the addition of the antibody. For each well, 0.5 µg of antibody was mixed with 1 µL of IncuCyte FabFluor-pH Antibody labeling reagent (Red) in 50 µL of T-cell medium. The mixture was then incubated at room temperature for 15 minutes and subsequently added to the cells. Antibody internalization kinetics were imaged and analyzed using the IncuCyte Live-Cell Analysis System. For higher-resolution imaging, confocal microscopy was performed using a ZEISS LSM 880; Hoechst 33342 was added to the cells prior to imaging to visualize nuclei.

### Cy5-labeling of tLNP-mRNA and internalization assay

To generate fluorescently labeled mRNA, 25% of the UTP in the *in vitro* transcription reaction was replaced with Cy5-labeled UTP, while all other steps of mRNA synthesis and purification were performed as described above. The resulting Cy5-labeled mRNA was then used to formulate tLNP–mRNA. For assessing tLNP internalization, activated T cells were seeded as described for the antibody internalization assay, and Hoechst 33342 was added to the culture medium to visualize nuclei. Subsequently, 0.5 µg of tLNP-Cy5-labeled mRNA encoding ZsGreen was added per well for confocal imaging of internalized mRNA and ZsGreen protein expression using a ZEISS LSM 880 confocal microscope. Each treatment condition included duplicate wells, and three fields of view per well were imaged for quantification.

### RNAseq analysis of tLNP-mRNA treated T cells

Donor-matched CD4⁺ or CD8⁺ T cells from three healthy donors were treated with aCD2- or aCD5-targeted LNP-ZsGreen mRNA at a dose of 1 μg mRNA per 1 × 10⁶ cells, with untreated cells and mouse IgG-conjugated LNP–ZsGreen mRNA serving as controls. After 24-hours, cells were collected, and total RNA was isolated using the Qiagen RNeasy Mini Kit. Sequencing data were processed in R (v4.2.2), and differential gene expression analysis was performed using DESeq2 (v1.38.3). Variance stabilizing transformation (VST) was applied for normalization, and batch effects were corrected using the removeBatchEffect function from the limma package (v3.54.2). Principal component analysis was visualized with ggplot2 [30]. Gene Set Enrichment Analysis (GSEA) was conducted to identify enriched pathways, with genes ranked using a metric that reflects both statistical significance and the directionality of change, calculated as log10(p-value) × sign(log2FoldChange). Gene Ontology (GO) enrichment and visualization were performed using the ClusterProfiler package (v4.6.2).

### *In vitro* generation of CAR T cells and killing assay

To evaluate the ability of aCD7/tLNP-mRNA to generate CAR T cells, mRNA encoding an anti-CD20 CAR construct was encapsulated in LNPs. T cells from two healthy donors were treated with 0.01, 0.1, or 1 μg of aCD7/tLNP-mRNA or with unconjugated LNP-mRNA dosage per million cells. After 24-hours, the cells were collected for flow cytometric analysis of CAR expression. To assess cytotoxic activity, CAR T cells were then co-cultured with CD20-expressing, luciferase-labeled Raji cells (ATCC, cat# CCL-86-LUC2) at effector-to-target ratios of 1:1, 2:1, 5:1, and 10:1. 10,000 Raji cells in 50 μL of Raji cell medium were seeded in 96-well optical-bottom plates that had been pretreated to enhance cell adherence, as described above, followed by a 2-hour incubation to promote attachment. Subsequently, CAR T cells in 50 μL of T-cell medium containing Incucyte® Cytotox Green Dye (for counting dead cells, Sartorius, Cat. 4633; 200 μM stock, diluted 1:2000) were added to each well. After 24-hours of co-culture, dead cells were imaged using an EVOS™ M5000 Imaging System (Thermo Fisher). Raji cell viability was then quantified by measuring luciferase activity using the ONE-Glo™ Luciferase Assay System (Promega, Cat# E6110) according to the manufacturer’s instructions.

### Humanized mouse model

All animal studies were conducted in accordance with the *Guide for the Care and Use of Laboratory Animals* (National Research Council). All procedures were approved by the Institutional Animal Care and Use Committee (IACUC) at the University of Pennsylvania, and the university’s animal facilities are accredited by the American Association for Accreditation of Laboratory Animal Care (AAALAC). PBMC-NSG humanized mice used for *in vivo* studies were obtained from the Jackson Laboratory (hu-PBMC-NSG, Cat. 745557). Female NSG™ mice (6–7 weeks old) were engrafted with human peripheral blood mononuclear cells (hu-PBMCs) by Jackson Laboratory. Peripheral blood was collected two weeks after engraftment to confirm successful humanization. Three weeks post-engraftment, humanized mice were randomized into treatment groups, and treated with tLNP-mRNA at a dosage of 2.5 μg mRNA per mouse. 24-hours after treatment, blood and spleen samples were collected to assess mRNA expression (ZsGreen reporter or anti-CD20 CAR construct) in T cells by flow cytometry.

### Immunohistochemistry staining

Spleens from aCD7/tLNP-aCD20 CAR-treated humanized mice were harvested and fixed in 4% paraformaldehyde (PFA) at 4 °C for 24-hours with orbital shaking at 100 rpm. Fixed tissues were washed in a graded ethanol series (30%, 50%, and 70%; each for 1-hour) before undergoing routine processing for paraffin embedding and sectioning at a thickness of 4 µm. For immunohistochemistry, antigen retrieval was performed in Tris-EDTA buffer (pH 9.0). Tissue sections were blocked with 10% goat serum for 15 minutes and incubated overnight at 4 °C with primary antibodies against CD20 (SP32, Abcam, Cat# ab64088; 1:100) and Granzyme B (D6E9W, Cell Signaling Technology, Cat. 46890; 1:100). After washing, secondary antibody incubation and chromogenic detection were performed using the Rabbit-Specific IHC Polymer Detection Kit, HRP/DAB (Abcam, Cat# ab209101), following the manufacturer’s instructions. Slides were counterstained with hematoxylin, mounted using an aqueous mounting medium (Abcam, Cat# ab264230), and imaged using an Aperio FL Scanning System (Leica).

### Statistical analysis

Data are presented as mean ± standard deviation (SD). Statistical differences between two groups were assessed using Student’s *t*-test, and comparisons among multiple groups were performed using one-way ANOVA. Significance thresholds were defined as: n.s. (not significant), \**p* < 0.05, \*\**p* < 0.01, and \*\*\**p* < 0.001. Statistical analysis was performed with GraphPad Prism 10.1.2.

## Results

### Comparison of targeting moieties for tLNP-mediated mRNA delivery efficiency to human primary T cells *in vitro*

To evaluate the relative performance of different targeting moieties systematically under uniform experimental conditions, we compared the mRNA delivery efficiency of LNPs conjugated with monoclonal antibodies against several highly expressed and T cell-specific surface markers: CD2, CD4, CD5, CD7, CD8, and CD4/8. Antibodies were covalently conjugated to LNP-mRNA via N-succinimidyl-S-acetylthioacetate (SATA)-maleimide chemistry (**Fig. 1A**, Methods) [20, 27]. Following antibody conjugation, the hydrodynamic diameter of LNPs increased from 94.4 nm to an average of 116.6 nm, as measured by dynamic light scattering. The zeta potential of tLNPs showed only minor changes relative to unmodified LNPs (**Fig. 1B**). To evaluate the impact of each targeting moiety on delivery efficiency of tLNP, we encapsulated a ZsGreen reporter mRNA as the payload. We treated activated human pan T cells at varying doses of each tLNP formulation (0.01, 0.1, or 1 μg mRNA per 100,000 cells) for 24-hours, followed by flow cytometric analysis of ZsGreen expression (**Fig. 1C**). Untreated cells or unmodified LNP-treated cells showed minimal ZsGreen signal in CD3⁺ T cells, whereas all antibody-conjugated tLNPs produced robust, dose-dependent reporter mRNA expression (**Fig. 1D, upper panel; Fig. 1E**). At the 1 μg dose, aCD2, aCD5, and aCD7/tLNPs achieved the highest efficiencies, with ∼80% of T cells expressing ZsGreen (aCD2: 80.6 ± 1.7%; aCD5: 83.3 ± 1.3%; aCD7: 76.4 ± 2.6%). In comparison, aCD4/tLNPs (target CD4) and aCD4/8-tLNPs (target both CD4 and CD8) yielded moderate efficiencies (∼60%), and aCD8/tLNPs were the least effective (17.0 ± 1.0%) (**Fig. 1D, E**).

**Fig. 1.**
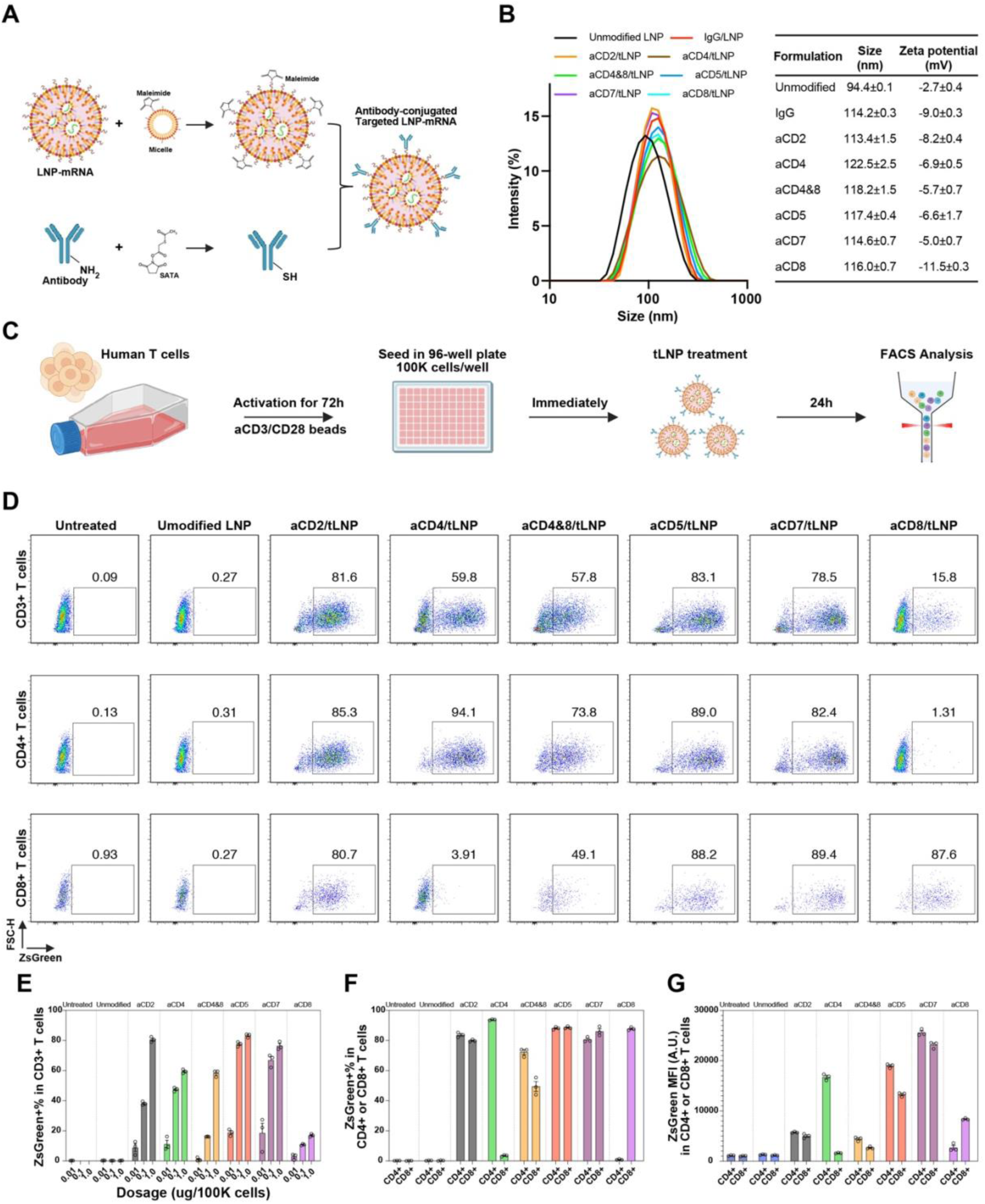
Comparison of targeting moieties for tLNP-mediated mRNA delivery efficiency to human primary T Cells *in vitro*. **(A)** Schematic of conjugating antibodies to the surface of LNP-mRNA through the SATA-maleimide chemistry. **(B)** Physical properties of LNP with vs. without antibody conjugation. **(C)** The analytic scheme of human T cells treated with tLNPs bearing targeting moieties against different T cell receptors. **(D)** The ZsGreen expression in CD3^+^, CD4^+^, and CD8^+^ T cells in each indicated group at 1 μg mRNA per 100,000 cells dosage, analyzed with flow cytometry. **(E)** ZsGreen+% in CD3^+^ T cells 24-hours after treated with each tLNP formulation at 0.01, 0.1, or 1 μg mRNA per 100,000 cells dosage. **(F)** ZsGreen+% in CD4^+^ VS. CD8^+^ T cells 24-hours after treated with each tLNP formulation at 1 μg mRNA per 100,000 cells dosage. **(G)** Mean fluorescence intensity of ZsGreen+ CD4^+^ VS. CD8^+^ T cells. *n* = 3 biological replicates. Data are represented as mean ± SD.

We next assessed subtype selectivity by quantifying ZsGreen expression in CD4⁺ and CD8⁺ T cell subsets (**Fig. 1D, middle and lower panels; Fig. 1F**). At 1 μg mRNA dose, aCD2, aCD5, and aCD7/tLNPs efficiently transfected both subsets, with comparable expression levels (aCD2: 83.7 ± 1.6% in CD4⁺ vs. 79.9 ± 0.7% in CD8⁺; aCD5: 88.1 ± 0.9% vs. 88.6 ± 0.8%; aCD7: 80.6 ± 1.6% vs. 86.1 ± 3.0%). In contrast, aCD4/tLNPs specifically transfected CD4⁺ T cells, while aCD8/tLNPs selectively targeted CD8⁺ T cells. The dual aCD4/8-tLNPs targeted both subsets but at lower efficiency than aCD2, aCD5, or aCD7-tLNPs formulations. To further compare the level of protein output, we quantified the mean fluorescence intensity of ZsGreen (**Fig. 1G**). Among the top-performing tLNPs, aCD7/tLNPs produced the strongest expression, with MFI values of 25,583 A.U. in CD4⁺ T cells and 23,131 A.U. in CD8⁺ T cells. aCD5/tLNPs yielded moderate expression (18,970 and 13,271 A.U., respectively), while aCD2/tLNPs showed lower intensity (5,742 and 5,008 A.U.). Together, these data identify aCD7/tLNPs as the most efficient formulation for mRNA delivery to human T cells, among the tLNPs tested in this study, combining broad subset targeting with the highest expression strength.

### Antibody-receptor internalization capacity, not receptor abundance, determines T-cell uptake of tLNP-mRNA

We next investigated factors influencing the mRNA delivery efficiency of tLNPs bearing different targeting moieties. We initially hypothesized that the abundance of target receptors on the T cell surface determines the efficiency of tLNP-mediated mRNA delivery. Under this hypothesis, the higher delivery efficiency observed with aCD7/tLNP compared to aCD2/tLNP would suggest greater CD7 expression than CD2. To examine this, we first assessed the expression levels of targeted receptors on T cells using publicly available datasets. Specifically, we referenced the Human Protein Atlas and the Schmiedel Immune Cell Dataset [31], which provide transcriptomic profiles of activated T cells from 91 healthy donors. Surprisingly, CD2 was expressed at the highest level among all targeted surface receptors, approximately fivefold higher than CD7 and CD5 (**Fig. 2A**). We further assessed the receptor abundance at the protein level using flow cytometry. Consistent with the trend in transcriptomic data, CD2 protein levels were approximately three times higher than those of CD5 and twice as high as those of CD7 (**Fig. 2B, C**). Counterintuitively, despite the higher abundance of CD2, aCD2/tLNP-ZsGreen transfected T cells yielded only one-third the ZsGreen MFI of aCD5/tLNP and one-fifth that of aCD7/tLNP, indicating that receptor abundance alone does not account for differences in tLNP-mRNA delivery efficiency and protein expression strength.

**Fig. 2.**
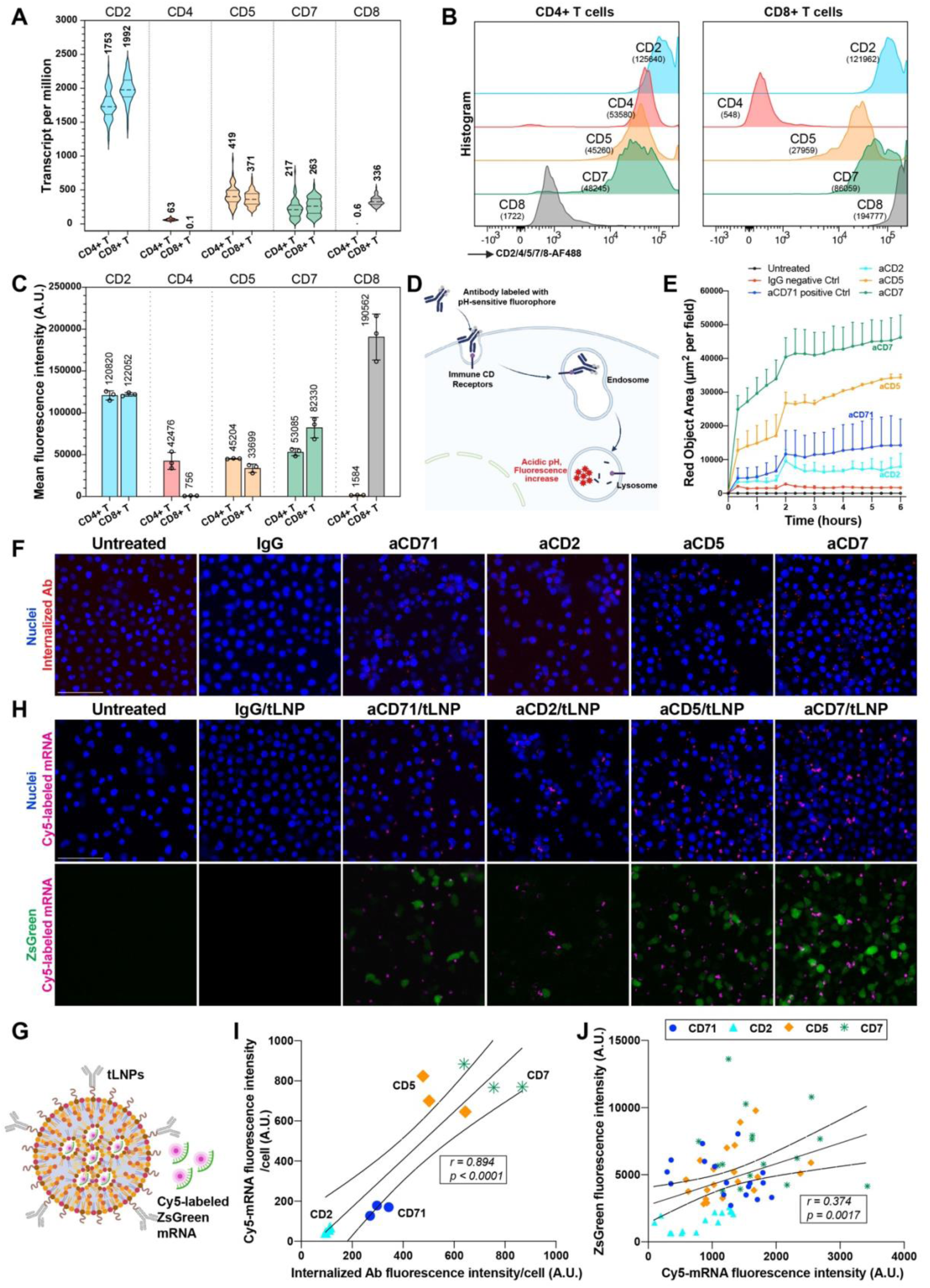
Higher antibody-receptor internalization, rather than receptor density, correlates with increased T-cell uptake of tLNP-mRNA. **(A)** Transcript levels of CD2, CD4, CD5, CD7, and CD8 in activated human CD4⁺ and CD8⁺ T cells analyzed using the Schmiedel Immune Cell Dataset (Human Protein Atlas). **(B)** Flow cytometry histograms showing protein expression of CD2, CD4, CD5, CD7, and CD8 in activated human CD4⁺ and CD8⁺ T cells. **(C)** Quantification of protein expression shown in (B). **(D)** Schematic of the IncuCyte® Fabfluor-pH antibody labeling system for quantitative analysis of antibody internalization via fluorescence. **(E)** Antibody internalization kinetics in activated T cells measured using the IncuCyte® Live-Cell Analysis System. **(F)** Representative images showing internalized antibodies (red) 4-hours after incubation with labeled antibodies. **(G)** Schematic showing Cy5-labeled mRNA encoding ZsGreen loaded into tLNPs to track mRNA uptake by T cells. **(H)** Representative images of internalized Cy5-labeled mRNA and ZsGreen expression 4-hours after tLNP treatment. **(I)** Correlation between antibody internalization (from F) and intracellular Cy5-mRNA accumulation (from H); each point represents one biological replicate. **(J)** Correlation between intracellular Cy5-mRNA and ZsGreen protein expression; each point represents a single cell, and cells from three biological replicates were pooled for plotting. *n* = 3 biological replicates for all experiments. Data are shown as mean ± SD. Scale bar = 100 μm.

We alternatively hypothesized that, rather than receptor density, the internalization of the antibody-receptor complex determines the efficiency of tLNP-mRNA uptake. To test this, we employed the IncuCyte® Fabfluor-pH antibody labeling system to quantitatively assess antibody internalization kinetics upon receptor binding. In this system, antibodies served as LNP targeting moieties are tagged with Fab fragments conjugated to pH-sensitive fluorophores, upon internalized into the cell and exposed to the acidic lysosomal environment (pH 4.5–5.5), the fluorophores exhibit a marked increase in fluorescence (**Fig. 2D**). Employing this labeling approach in combination with the IncuCyte® Live-Cell Analysis System, we monitored the internalization of antibodies targeting CD2, CD5, and CD7 in activated human T cells. IgG served as a negative control, while CD71 (transferrin receptor, known to undergo extensive receptor cycling but not T-cell-specific) was included as a positive control [32, 33]. We found that aCD7 exhibited the highest level of antibody internalization, followed by aCD5, both of which exceeded the positive control aCD71 (**Fig. 2E**). In contrast, aCD2 showed substantially lower internalization. Confocal imaging at 4 hours after antibody addition further confirmed the higher internalization and lysosomal accumulation of aCD7 (**Fig. 2F**). To assess whether the higher internalization of aCD7 relative to aCD2 is antibody-clone dependent, we evaluated three distinct clones of aCD7 and aCD2 antibodies and compared their internalization kinetics. The capacity of these antibodies to bind to their targeting receptors was confirmed by flow cytometry **(Fig. S1A, B)**. Using the FabFluor-pH labeling system, we found that all aCD7 clones exhibited substantially higher internalization than the aCD2 clones, averaging a four times increase (aCD7: 66873 ± 10075 A.U. VS. aCD2: 17329 ± 7052 A.U.), despite minor clone-to-clone variations **(Fig. S1C, D)**. These results indicate that aCD7 internalizes more efficiently than aCD2 upon receptor binding, largely regardless of the antibody clone.

To further determine whether the higher internalization of aCD7 antibodies correlates with increased uptake of aCD7/tLNPs, we encapsulated Cy5-labeled mRNA encoding ZsGreen within tLNPs (**Fig. 2G**). Human T cells were treated with aCD2-, aCD5-, or aCD7-conjugated tLNPs, while IgG-tLNP and aCD71-tLNP served as negative and positive controls, respectively. 4-hours post-transfection, confocal imaging revealed the greatest accumulation of Cy5-labeled mRNA in cells treated with aCD7/tLNP, followed by aCD5/tLNP, aCD71/tLNP, and finally aCD2/tLNP, a pattern consistent with the antibody internalization kinetics (**Fig. 2H upper panel)**. Quantitative analysis showed a strong positive correlation between the level of antibody internalization shown in Fig. 3F and the intracellular accumulation of Cy5-mRNA shown in Fig. 3H (**Pearson *r* = 0.894, \*\**p* < 0.0001) (**Fig. 2I**), indicating that antibody internalization strongly corelates tLNP-mRNA uptake efficiency. Further analysis of the ZsGreen protein level in each cell revealed a significant, although weaker, positive correlation to the amount of Cy5-mRNA uptake (\*\**r* = 0.374, *p* = 0.0017) (**Fig. 2J**). The reduced correlation may reflect differences in mRNA translation activity in T cells following distinct tLNP treatments. Consistent with this hypothesis, RNA sequencing and gene set enrichment analysis comparing aCD7/tLNP-ZsGreen treated T cells compared with aCD2/tLNP-ZsGreen treated T cells, revealed strong enrichment of pathways related to protein translation, ribosome biogenesis, and mRNA metabolism in CD4⁺ T cells **(Fig. S2A, B, C)**. In CD8⁺ T cells, aCD7/tLNP treatment showed positive, though more modest, enrichment for mRNA processing and translation accompanied by significant downregulation of mRNA decay pathways compared with aCD2/tLNP treatment **(Fig. S2D, E)**. These findings indicate that aCD7/tLNP-treated T cells may be in a more active state for mRNA translation than those treated with aCD2/tLNP.

**Fig. 3.**
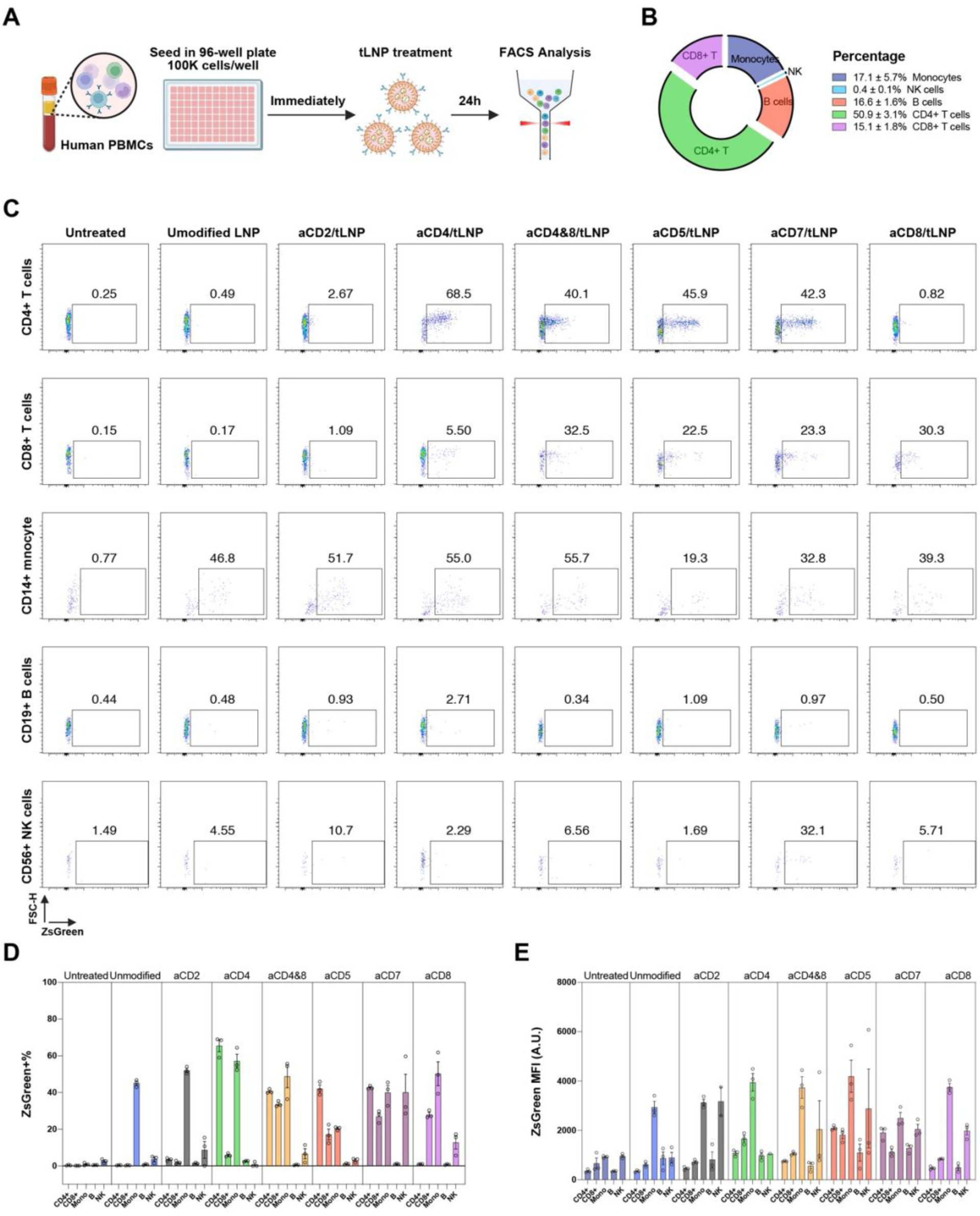
aCD7/tLNPs efficiently deliver mRNA to T cells in the presence of other immune cells. **(A)** Experimental scheme of human PBMCs treated with tLNPs bearing targeting moieties against different T cell surface receptors. **(B)** Immune cell composition of PBMCs from three healthy donors used for tLNP treatment. **(C)** ZsGreen expression in CD4⁺ and CD8⁺ T cells, CD14⁺ monocytes, CD19⁺ B cells, and CD56⁺ NK cells 24-hours after treatment with each tLNP formulation at 1 μg mRNA per 100,000 cells dosage, analyzed by flow cytometry. **(D)** Quantification of ZsGreen⁺ cell percentages in each immune cell subtype. **(E)** Mean fluorescence intensity of ZsGreen⁺ cells in each immune cell subtype. *n* = 3 biological replicates. Data are shown as mean ± SD.

In summary, our results demonstrate that the capacity of antibody-receptor internalization, rather than receptor abundance, is the key determinant of tLNP efficiency for mRNA delivery to human T cells. This internalization capacity appears to be an intrinsic property of each receptor and largely independent of the binding antibody clone.

### aCD7/tLNP efficiently delivers mRNA to T cells in the presence of other immune cells

To evaluate whether aCD7/tLNPs can efficiently deliver mRNA to T cells in the presence of other immune cell populations, we assessed their performance using human peripheral blood mononuclear cells (PBMCs) (**Fig. 3A**), which more closely recapitulate the cellular complexity *in vivo*. We first characterized the immune cell composition of PBMCs from three healthy donors and found that they comprised CD4⁺ T cells (50.9 ± 3.1%), CD8⁺ T cells (15.1 ± 1.8%), B cells (16.6 ± 1.6%), monocytes (17.1 ± 5.7%), and NK cells (0.4 ± 0.1%) (**Fig. 3B**). We then treated the PBMCs with tLNPs bearing targeting moieties against CD2, CD4, CD5, CD7, CD8, or CD4/8, each loaded with ZsGreen mRNA as a reporter. 24-hours post-treatment, we performed flow cytometry to quantify the percentage of ZsGreen⁺ cells and MFI within each immune subset, with a focus on T cells. Even in the presence of other immune cells, aCD7/tLNPs maintained the highest T-cell targeting efficiency (42.7 ± 0.9% in CD4⁺ T cells and 27.0 ± 3.4% in CD8⁺ T cells), followed by aCD5/tLNPs (42.1 ± 3.6% in CD4⁺ T cells and 17.1 ± 5.1% in CD8⁺ T cells) (**Fig. 3C, D**). The absolute efficiencies were approximately half of those observed in pure T cells shown in Fig. 1E, likely due to competition for nanoparticle uptake among different cell types, particularly monocytes, which exhibited consistently high ZsGreen⁺ percentages across all treatment conditions (**Fig. 3C, D**). Among other PBMC subsets, B cells showed negligible ZsGreen expression, whereas NK cells displayed elevated ZsGreen⁺ levels specifically under aCD7/tLNP treatment, consistent with the high CD7 expression on NK cells [34, 35]. In terms of mRNA expression strength, aCD7/tLNPs again ranked among the most effective formulations, with average MFIs of 1,907 A.U. in CD4⁺ T cells and 1,124 A.U. in CD8⁺ T cells (**Fig. 3E**).

Notably, these MFI values were 10–20 times lower than those observed in purified, activated T-cell cultures shown in Fig. 1G. This reduction likely reflects both decreased tLNP bioavailability due to competition from other cell types and the absence of the *in vitro* T-cell activation step in PBMC experiments, which was omitted to preserve other immune populations, as monocytes and NK cells do not survive the 72-hour activation process **(Fig. S3A, B)**. When comparing the performance of aCD7/tLNP-ZsGreen in activated versus non-activated T cells, we found that T-cell activation substantially enhanced transfection efficiency, with the ZsGreen⁺ fraction approximately doubling in both CD4⁺ and CD8⁺ T cells **(Fig. S3C–F)**. Likewise, ZsGreen MFI increased markedly upon activation, from 1,541 to 8,649 in CD4⁺ T cells and from 1,807 to 16,940 in CD8⁺ T cells **(Fig. S3G)**. Together, these results demonstrate that aCD7/tLNPs efficiently deliver mRNA to T cells even in the presence of other immune cell populations, and that T-cell activation further enhances mRNA expression levels.

### aCD7/tLNP efficiently delivers mRNA to T cells *in vivo*

To evaluate whether aCD7/tLNPs can achieve efficient mRNA delivery to T cells *in vivo*, we employed a humanized mouse model generated by engrafting human PBMCs into NOD.Cg-*Prkdc^scid^ Il2rg^tm1Wjl^*/SzJ (NSG) mice, a well-established system for studying human T-cell biology *in vivo* [36–38]. We first characterized the kinetics of human immune cell reconstitution in this model. By four weeks post-engraftment, human CD45⁺ cells constituted approximately 50% of splenocytes and 20% of circulating white blood cells, the majority of which were human CD4⁺ or CD8⁺ T cells **(Fig. S4)**, indicating a robust level of humanization. At this time point, mice were administered either unmodified LNP-ZsGreen or aCD7/tLNP-ZsGreen via retro-orbital injection at a dose of 2.5 µg/mouse. 24-hours post-injection, spleen and blood samples were collected to assess ZsGreen expression in human T cells (**Fig. 4A**). In splenic T cells, aCD7/tLNP treatment resulted in 59.3% ZsGreen⁺ CD4⁺ T cells and 88.7% ZsGreen⁺ CD8⁺ T cells (**Fig. 4B-D**). T cells isolated from peripheral blood displayed ZsGreen⁺ frequencies and MFI values comparable to those in splenic T cells (**Fig. 4E–G**). Notably, in both spleen and blood, CD8⁺ T cells consistently exhibited higher ZsGreen⁺ percentages and fluorescence intensities than CD4⁺ T cells, for example, 88.7% vs. 59.3% ZsGreen⁺ and 20,912 A.U. vs. 7,028 A.U. MFI in the spleen, likely reflecting the considerably higher CD7 expression on CD8⁺ than CD4⁺ T cells, as shown in fig. 2B, C by flow cytometry. Collectively, these findings demonstrate that aCD7/tLNPs efficiently delivers mRNA to human T cells *in vivo*, with preferential uptake and expression in CD8⁺ T cells compared to CD4⁺ T cells.

**Fig. 4.**
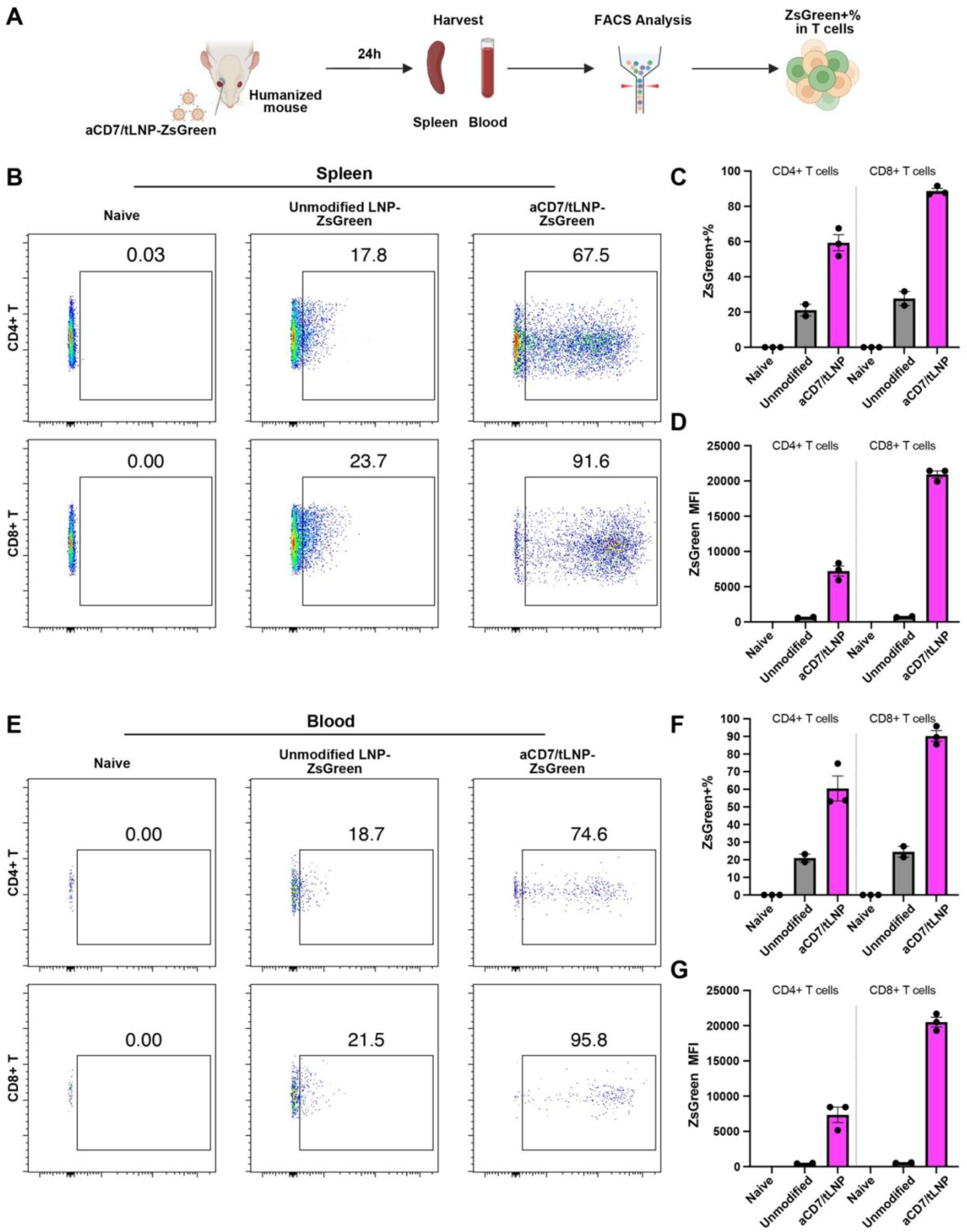
aCD7/tLNP efficiently delivers mRNA to T cells *in vivo*. **(A)** Experimental scheme for evaluating aCD7/tLNP-mediated mRNA delivery to human T cells *in vivo*. **(B)** ZsGreen expression in human CD4⁺ and CD8⁺ T cells in the spleen of humanized mice 24-hours after treatment with each tLNP formulation (2.5 μg mRNA per mouse). **(C)** Quantification of ZsGreen⁺ cell percentages in human CD4⁺ and CD8⁺ T cells in the spleen. **(D)** Mean fluorescence intensity of ZsGreen⁺ CD4⁺ and CD8⁺ T cells in the spleen. **(E)** ZsGreen expression in human CD4⁺ and CD8⁺ T cells in peripheral blood. **(F)** Quantification of ZsGreen⁺ cell percentages in human CD4⁺ and CD8⁺ T cells in the blood. **(G)** MFI of ZsGreen⁺ CD4⁺ and CD8⁺ T cells in the blood. *n* = 3 mice for naïve and aCD7/tLNP-ZsGreen–treated groups; *n* = 2 for the unmodified LNP-ZsGreen control group. Data are shown as mean ± SD.

### aCD7/tLNP loaded with anti-CD20 CAR mRNA efficiently generates functional CAR T cells *in vitro*

To evaluate the potential of aCD7/tLNPs for engineering human T cells into CAR T cells, we loaded the tLNP with mRNA encoding anti-CD20 CAR. Activated T cells from two different donors were treated with a range of doses, 0.01, 0.1, and 1 µg of aCD7/tLNPs-anti-CD20 CAR mRNA per million cells. 24-hours post-treatment, cells were collected and stained for the G4S linker within the anti-CD20 CAR construct to quantify CAR expression. We found that, while treatment with unmodified LNPs resulted in minimal CAR expression across all doses (**Fig. 5A, upper panel; Fig. 5B, C**), aCD7/tLNP-treated cells displayed a clear dose-dependent increase in the proportion of CAR⁺ cells within both CD4⁺ and CD8⁺ T-cell subsets. At the highest dose (1 µg mRNA/million cells), donor ND615 exhibited 48.1% CAR⁺ CD4⁺ T cells and 76.0% CAR⁺ CD8⁺ T cells, while a second donor showed 63.3% CAR⁺ CD4⁺ T cells and 64.0% CAR⁺ CD8⁺ T cells (**Fig. 5A, lower panel; Fig. 5D, E**). These results demonstrate that aCD7/tLNPs enable highly efficient mRNA delivery for the generation of CAR T cells *in vitro*.

**Fig. 5.**
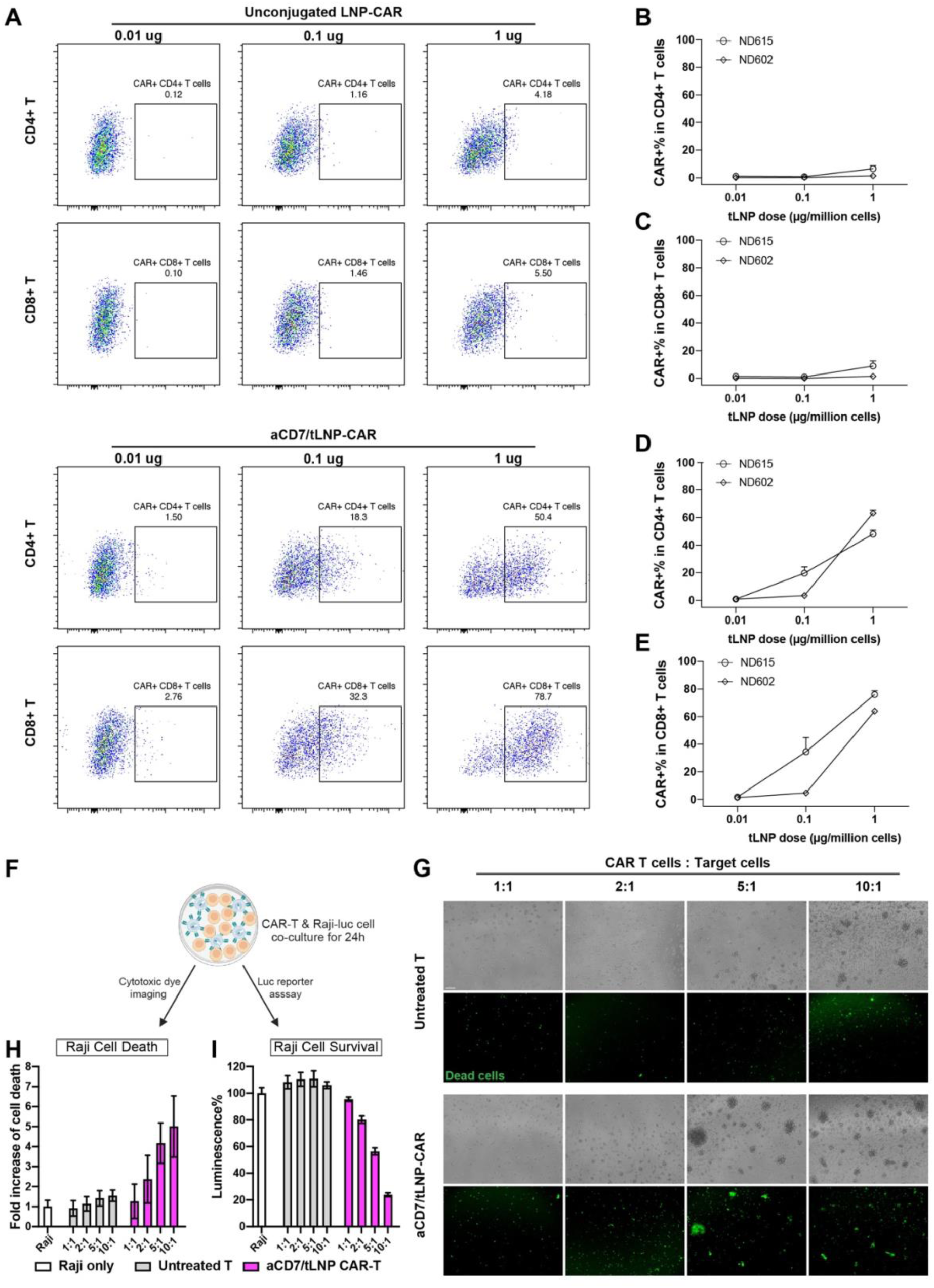
aCD7/tLNP loaded with anti-CD20 CAR mRNA efficiently generates functional CAR T cells *in vitro*. **(A)** CAR expression in activated human T cells 24-hours after treatment with unconjugated LNP–CAR mRNA or aCD7/tLNP-CAR mRNA at doses of 0.01, 0.1, or 1 μg mRNA per million cells. **(B and C)** Quantification of CAR expression in CD4⁺ and CD8⁺ T cells treated with unconjugated LNP-CAR mRNA. **(D and E)** Quantification of CAR expression in CD4⁺ and CD8⁺ T cells treated with aCD7/tLNP-CAR mRNA. **(F)** Experimental scheme for evaluating the cytolytic activity of CAR T cells against target cells. **(G)** Representative images showing Cytotox Green fluorescence indicating Raji cell death in co-cultures with T cells treated with unmodified LNP-CAR or aCD7/tLNP-CAR at effector-to-target (E: T) ratios of 1:1, 2:1, 5:1, and 10:1. **(H)** Quantification of Cytotox Green fluorescence shown in (G). **(I)** Quantification of Raji cell viability in co-culture based on luciferase activity. *n* = 2 biological replicates. Data are shown as mean ± SD. Scale bar = 100 μm.

We next assessed whether the CAR T cells generated with aCD7/tLNPs were functionally capable of killing their target cells. We co-cultured CAR T cells with the lymphoblast-like Raji cell line, which was verified to express the CD20 antigen.**(Fig. S5)**. CAR T cells were co-cultured with Raji cells at effector-to-target (E: T) ratios of 1:1, 2:1, 5:1, and 10:1. To visualize target cell killing, we added Cytotox Green dye, which fluoresces upon binding to the DNA of dead cells, and imaged cultures after 24-hours. T cells treated with unmodified LNPs exhibited minimal cytotoxic activity across all E: T ratios. In contrast, aCD7/tLNP-engineered CAR T cells showed a prominent ratio-dependent increase in Raji cell killing, with ∼5-fold higher Cytotox Green fluorescence at 5:1 and 10:1 ratios compared to Raji-only controls (**Fig. 5F-H**). We also quantified cytotoxicity and Raji cell survival by measuring luciferase activity, as the Raji cell line expresses a luciferase reporter. Consistent with imaging results, co-culture with aCD7/tLNP-generated CAR T cells led to a dose-dependent reduction in luciferase signal, confirming effective target cell killing (**Fig. 5I**). Collectively, these findings demonstrate that the aCD7/tLNP platform efficiently delivers CAR mRNA into human T cells, generating potent and functional CAR T cells capable of antigen-specific cytotoxicity *in vitro*.

### aCD7/tLNP loaded with anti-CD20 CAR mRNA efficiently generates CAR T cells *in vivo* and depletes B cells

To further evaluate the ability of aCD7/tLNPs to deliver anti-CD20 CAR mRNA *in vivo* for CAR T-cell generation and B-cell depletion, we employed the PBMC-NSG humanized mouse model. Mice were administered either aCD7/tLNP-ZsGreen or aCD7/tLNP-CAR at 2.5 µg mRNA via retro-orbital injection. 24-hours post-injection, spleens were harvested and divided for analysis: one half for flow cytometry to assess CAR expression efficiency, and the other for immunohistochemistry (IHC) to evaluate depletion of CD20⁺ B cells (**Fig. 6A**). In aCD7/tLNP-CAR-treated mice, we found an average of 33.3% of human CD4⁺ T cells and 50.6% of CD8⁺ T cells expressed the CAR construct, whereas CAR expression was minimal in both untreated controls and aCD7/tLNP-ZsGreen–treated mice (**Fig. 6B–D**). Consistent with our *in vitro* findings that aCD7/tLNP mediates more efficient mRNA delivery to T cells than aCD2/tLNP, humanized mice treated with aCD2/tLNP-CAR exhibited substantially lower CAR expression, averaging 11.5% in CD4⁺ T cells and 20.2% in CD8⁺ T cells **(Fig. S6)**. To determine whether the *in vivo*-generated CAR T cells were functionally active, we quantified CD20⁺ B cells in the spleen. In untreated mice (*n* = 6) and aCD7/tLNP-ZsGreen–treated control mice, CD20⁺ B cells accounted for 4.1% and 7.5% of splenocytes, respectively. In contrast, aCD7/tLNP-CAR-treated mice exhibited only 0.5% CD20⁺ B cells, representing a significant reduction and indicating potent CAR T-cell-mediated B-cell clearance (**Fig. 6B, lower panel; Fig. 6E**). IHC staining further confirmed near-complete B-cell depletion in the spleen of aCD7/tLNP-CAR–treated mice. Compared with approximately 500 CD20⁺ B cells per field observed in naïve and aCD7/tLNP-ZsGreen–treated control animals, fewer than 20 CD20⁺ cells per field were detected following aCD7/tLNP-CAR treatment (**Fig. 6F, upper panel; Fig. 6G**). Moreover, strong Granzyme B staining was observed in the spleens of aCD7/tLNP-CAR-treated mice, but not in controls (**Fig. 6F, lower panel)**, indicating robust cytotoxic activity of the generated CAR T cells. Collectively, these findings demonstrate that aCD7/tLNPs loaded with anti-CD20 CAR mRNA efficiently generate functional CAR T cells *in vivo*, leading to potent B-cell depletion.

**Fig. 6.**
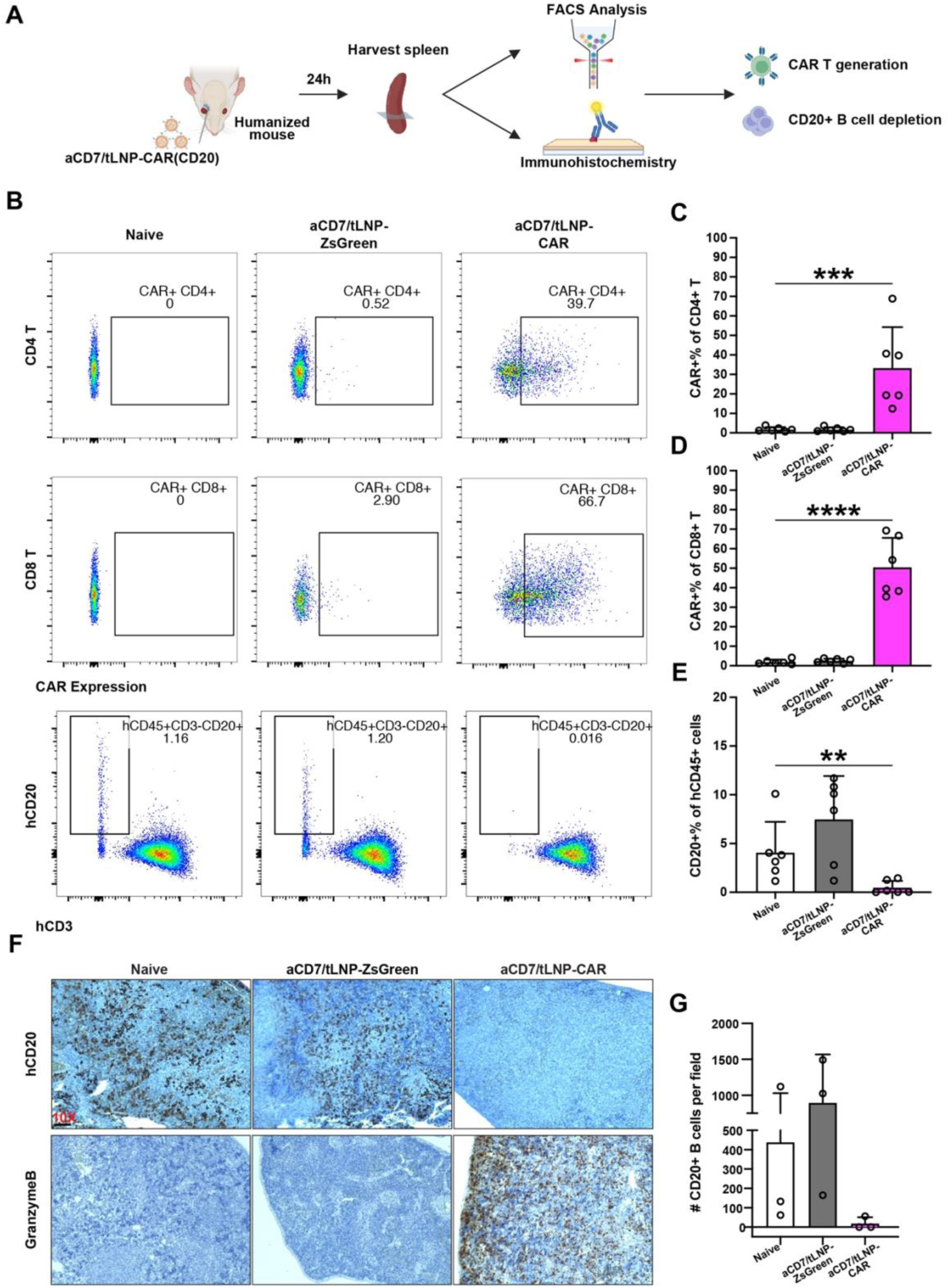
aCD7/tLNP loaded with anti-CD20 CAR mRNA efficiently generates CAR T cells *in vivo* and depletes B cells. **(A)** Experimental scheme for evaluating aCD7/tLNP-mediated CAR T-cell generation *in vivo* and their cytolytic activity against target cells. **(B)** CAR expression in CD4⁺ and CD8⁺ T cells in the spleen and levels of remaining CD20⁺ B cells 24-hours after treatment with aCD7/tLNP-ZsGreen (control) or aCD7/tLNP-CAR mRNA (2.5 μg per mouse), analyzed by flow cytometry. **(C and D)** Quantification of CAR expression in CD4⁺ and CD8⁺ T cells. **(E)** Quantification of CD20⁺ B cells in the spleen. **(F)** Representative immunohistochemistry images showing CD20⁺ B cells (upper panel) and Granzyme B (lower panel) in spleens from aCD7/tLNP-ZsGreen- or aCD7/tLNP-CAR-treated humanized mice. **(G)** Quantification of CD20⁺ B-cell density shown in (F). *n* = 6 mice for (B-E); *n* = 3 mice for (F and G). Data are shown as mean ± SD. Scale bar, 100 μm.

## Discussion

Antibody-functionalized tLNPs have emerged as a powerful platform for *in vivo* mRNA delivery to T cells and for generating CAR T cells directly *in vivo* [19]. Although multiple T-cell surface receptors, including CD4, CD5, CD7, and CD8, have been targeted by antibody-functionalized tLNPs [18, 20–22], the relative performance of tLNPs targeting these receptors for mRNA delivery to T cells under identical conditions and the mechanisms underlying their varied efficiencies remain unclear. Clarifying this question is essential for designing optimal tLNPs for *in vivo* CAR T-cell therapy. In this study, we systematically compared tLNPs targeting CD2, CD4, CD5, CD7, CD8, and CD4/8 to determine which receptor allows for the most efficient delivery of mRNA to T cells and the reasons behind this efficiency. We showed that aCD7/tLNPs consistently outperformed other targeting moieties, achieving the highest mRNA delivery *in vitro* and generating potent CAR T cells *in vivo*. By integrating measurements of receptor abundance, antibody–receptor internalization across multiple different antibody clones, and intracellular mRNA and protein output, we showed that although CD2 is the most abundant receptor on human T cell surface among the list of receptors targeted, it internalizes poorly and supports limited tLNP-mRNA uptake and protein expression. In contrast, antibody binding to CD7 undergoes rapid, robust, and clone-independent internalization, enabling superior tLNP-mRNA uptake and protein production. These results establish antibody-receptor internalization capacity, a property intrinsic to each receptor and largely independent of antibody clone, as the key parameter governing effective tLNP-mediated mRNA delivery to T cells and highlight CD7 as an optimal targeting receptor for *in vivo* CAR T-cell engineering.

Our *in vivo* studies using humanized mice showed that aCD7/tLNP-CAR generated substantially higher CAR expression than aCD2/tLNP–CAR, with an average of 33.3% versus 11.5% CAR⁺ in CD4⁺ T cells and 50.6% versus 20.2% CAR⁺ in CD8⁺ T cells (**Fig. 6, S6**). These results are consistent with our *in vitro* findings that antibody binding to CD7 internalizes far more efficiently than those binding to CD2 (**Fig. 2, S1**) and drives higher tLNP-mRNA delivery and ZsGreen reporter expression (**Fig. 1**). Notably, the *in vivo* transfection efficiency to generate CD8⁺ CAR T cells we observed at 24-hours post aCD7/tLNP-CAR treatment is nearly double that reported by Hunter *et al.* using aCD8/tLNP-CAR, despite our using a dose of only 2.5 μg mRNA compared with their 30 μg per mouse, over a tenfold higher amount [18]. Similarly, compared with the anti-FAP CAR generated using aCD5/tLNPs [22], our aCD7/tLNP still achieved nearly double the CAR⁺ percentage while using only one-quarter of the mRNA dose. Together, these data indicate that CD7-targeted tLNP–mRNA enables superior efficiency for in vivo CAR T-cell generation. We acknowledge, however, that these studies used different CAR constructs; thus, differences in the percentage of CAR⁺ cells may reflect not only variations in mRNA delivery efficiency but also differences in the translational efficiency of the respective CAR-encoding mRNAs. A direct head-to-head comparison of all targeting moieties *in vivo* under identical conditions will be important for definitive benchmarking in the future. An additional advantage of targeting CD7 is its high expression on NK cells [34, 35], which enables the simultaneous generation of CAR⁺ T cells and CAR⁺ NK cells, potentially enhancing target-cell clearance [35, 39]. This NK-cell transfection capacity is also reflected in our PBMC experiments, where aCD7/tLNP–ZsGreen yielded ∼30% ZsGreen⁺ NK cells, whereas other targeting moieties showed minimal NK-cell transfection (**Fig. 3**). Overall, CD7 represents a highly potent targeting receptor for *in vivo* CAR generation using tLNP-mRNA.

To our surprise, although antibody-receptor internalization is strongly associated with tLNP-mRNA uptake levels, the correlation between the levels of intracellular mRNA accumulation and protein output is more modest (**Fig. 2I, J**). One potential explanation for the partial uncoupling between intracellular mRNA levels and protein output is that engagement of different antibody–receptor pairs (e.g., aCD2–CD2 versus aCD7–CD7) may differentially modulate the translational activity of T cells. Antibody binding has been reported to exert agonistic effects and activate downstream signaling pathways [40, 41], and CD7 itself functions as a co-stimulatory molecule in T cells [42, 43]. Consistent with this possibility, our transcriptomic profiling revealed that aCD7/tLNP-treated T cells upregulate pathways associated with mRNA metabolism, translation, and ribosome biogenesis (**Fig. S2**). However, these observations are preliminary; whether aCD7/tLNPs truly prime T cells toward a more translation-active state remains a hypothesis and will require more comprehensive mechanistic investigation in the future.

Our study also highlights opportunities for further refinement. First, our selection of targeting moieties focused on receptors previously explored in *in vivo* CAR studies and therefore did not encompass all potential candidates. Incorporating additional T-cell–specific receptors—such as TCR, CD28, CCR7, and CXCR4—could further expand and strengthen the targeting repertoire [44–46]. Our finding that antibody-receptor internalization capacity dictates mRNA delivery efficiency provides a guiding principle for prioritizing new candidates and assessing whether any may even outperform aCD7/tLNP. Second, our framework identifying rapidly internalizing receptors as optimal targets for efficient tLNP-based mRNA delivery is derived from human T-cell data, and whether this principle extends to other cell types remains to be determined. Future studies will evaluate this internalization-based selection strategy in additional hard-to-transfect cell populations. Third, the PBMC-NSG humanized mouse model used in our *in vivo* studies provides robust human T-cell reconstitution but lacks other critical immune components, including NK cells and myeloid populations, and therefore does not fully recapitulate the complexity of the human immune system. Evaluating aCD7/tLNPs in models with more complete immune repertoires will be needed to more accurately assess performance and translational potential.

In summary, our study establishes antibody-receptor internalization capacity as a key design principle for selecting targeting moieties for tLNP-mediated mRNA delivery to T cells *in vivo*. We identify CD7 as a particularly effective receptor to target, enabling robust mRNA delivery to T cells and generation of *in vivo* CAR T cells. These findings provide a framework for rational selection of targeting receptors to optimize tLNP-mRNA-based *in vivo* CAR T-cell therapies and other T-cell engineering applications.

## CRediT authorship contribution statement

**Jianhao Zeng:** Conceptualization, Data curation, Formal analysis, Investigation, Methodology, Visualization, Writing – original draft, Writing – review and editing. **Tyler Ellis Papp:** Conceptualization, Data curation, Investigation, Methodology, Writing – review and editing. **Awurama Akyianu:** Data curation, Investigation, Methodology, Writing – review and editing. **Alejandra Bahena:** Investigation, Methodology, Writing – review and editing. **Lanfranco Leo:** Investigation, Methodology, Writing – review and editing. **Faris Halilovic:** Investigation, Methodology, Writing – review and editing. **Hamideh Parhiz:** Conceptualization, Funding acquisition, Supervision, Writing – review and editing

## Acknowledgments

We thank Max Eldabbas, Emileigh Maddox, Tanishk Sinha, and Jiayi Shu of the Human Immunology Core for their assistance with human T-cell collection. This core is supported in part by NIH grants P30 AI045008 and P30 CA016520. Flow cytometry data were generated at the Penn Cytomics and Cell Sorting Shared Resource Laboratory (RRID:SCR_022376), which is partially supported by the Abramson Cancer Center NCI Grant (P30 CA016520). We also acknowledge the Center for Molecular Studies in Digestive and Liver Diseases (P30 DK050306) and its Molecular Pathology and Imaging Core (RRID:SCR_022420) for histopathology support, as well as the Cell & Developmental Biology Microscopy Core (RRID:SCR_022373) for imaging assistance. We thank Linhui Chen and Taehyong Kim of the Bioinformatics Core at Penn for RNAseq data analysis. All these cores are affiliated with the University of Pennsylvania.

## Data and materials availability

Data are included within the paper and its supplemental information. Raw data collected are available from the corresponding author upon request.

