## Supplemental figures for "Rapid receptor internalization potentiates CD7-targeted lipid nanoparticles for efficient mRNA delivery to T cells and *in vivo* CAR T-cell engineering"

Jianhao Zeng *et al.*

**This file includes:**

Figs. S1 to S6

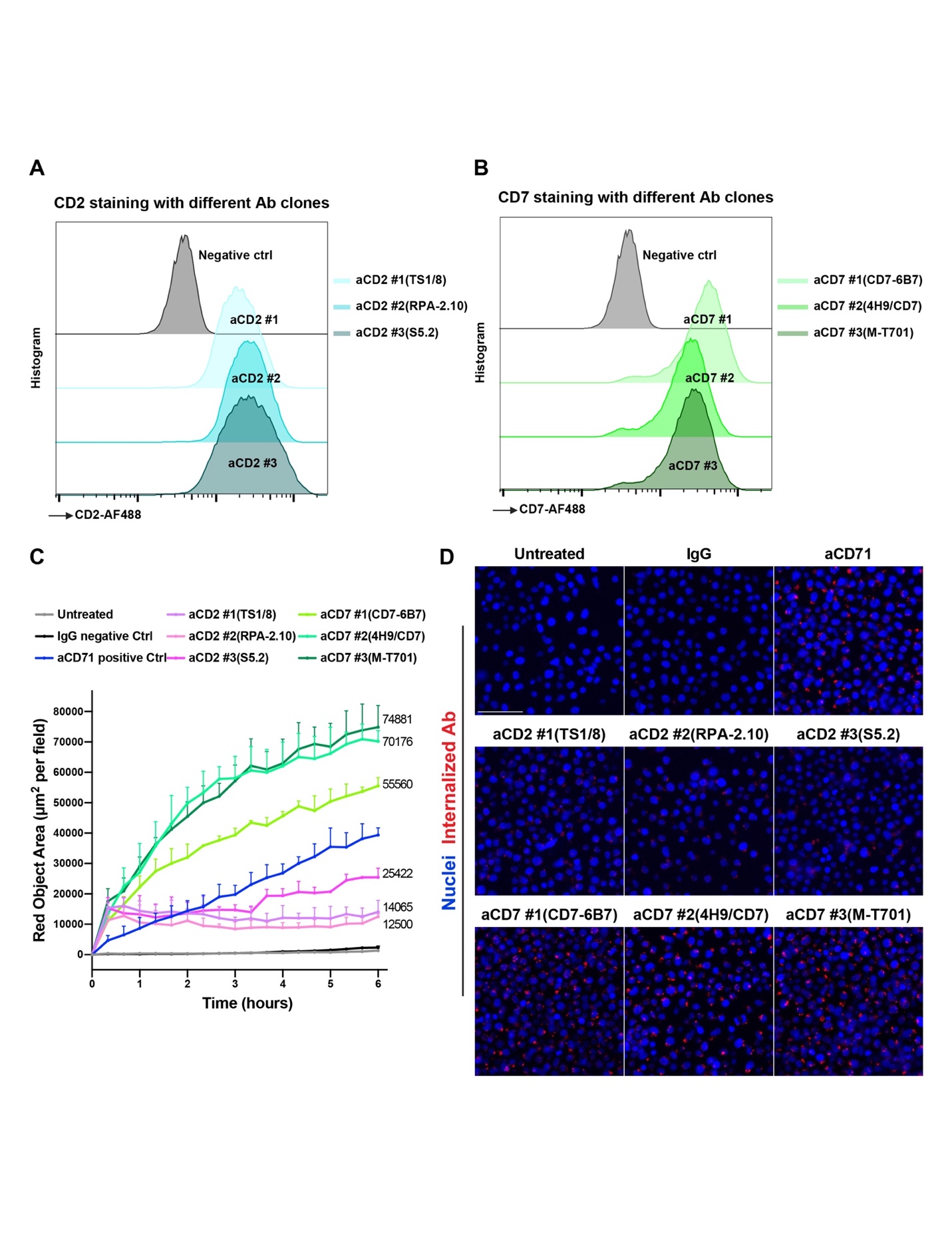

**Fig. S1. Higher antibody–receptor internalization of aCD7 than aCD2 across all tested antibody clones. (A-B)** Flow cytometric staining of primary human T cells using three monoclonal antibody clones targeting CD2 (A) or CD7 (B). Histograms show relative surface staining intensity compared to the isotype control. **(C)** Kinetics of antibody internalization for each CD2- and CD7-targeting clone, analyzed and quantified using the IncuCyte® FabFluor-pH antibody labeling system and the IncuCyte® Live-Cell Analysis System. **(D)** Representative images of internalized antibodies (red) 12-hours after incubation with the indicated labeled clones. *n* = 3 biological replicates for all experiments. Data represent mean ± SD. Scale bar, 100 μm.

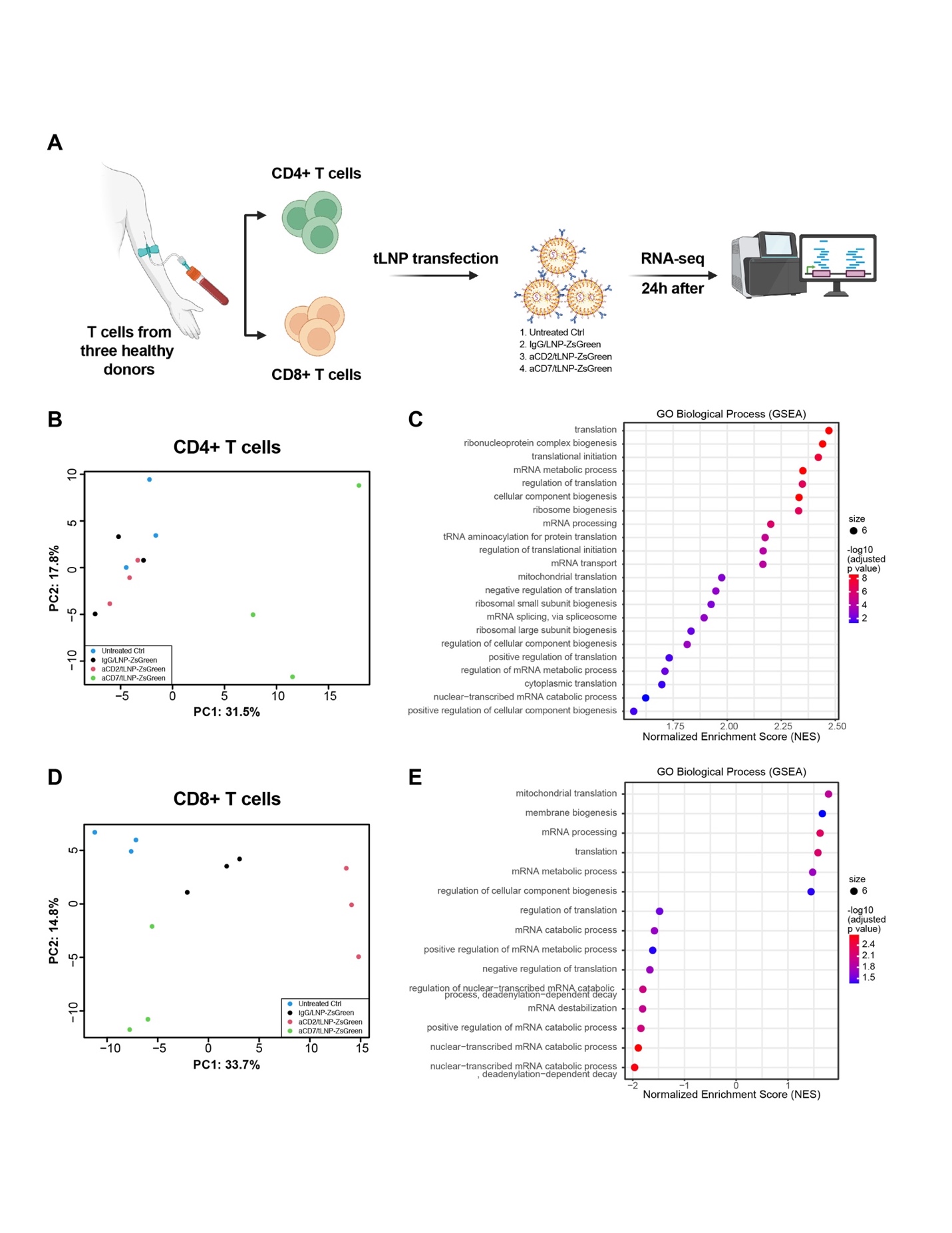

**Fig. S2. Transcriptomic profiling of CD4⁺ and CD8⁺ T cells following aCD2- or aCD7-tLNP-mediated mRNA delivery.** **(A)** Schematic overview of the experimental workflow. CD4⁺ and CD8⁺ T cells from three healthy donors were transfected with the indicated formulations, untreated control, IgG/tLNP-ZsGreen, aCD2/tLNP-ZsGreen, or aCD7/tLNP-ZsGreen, at a dose of 1 μg mRNA per 1 × 10⁶ cells. Total mRNA was extracted 24-hours after transfection for RNA sequencing. **(B-C)** Principal component analysis (B) and Gene Ontology (GO) biological process gene set enrichment analysis (GSEA) (**C**) for CD4⁺ T cells, comparing aCD7/tLNP-treated cells with aCD2/tLNP-treated cells. Selected pathways related to mRNA metabolism, translation, and ribosome biogenesis are shown. PCA (**D**) and GO biological process GSEA (**E**) for CD8⁺ T cells, comparing aCD7/tLNP-treated cells with aCD2/tLNP-treated cells, with the same selected categories of pathways displayed to enable parallel comparison across T-cell subsets.

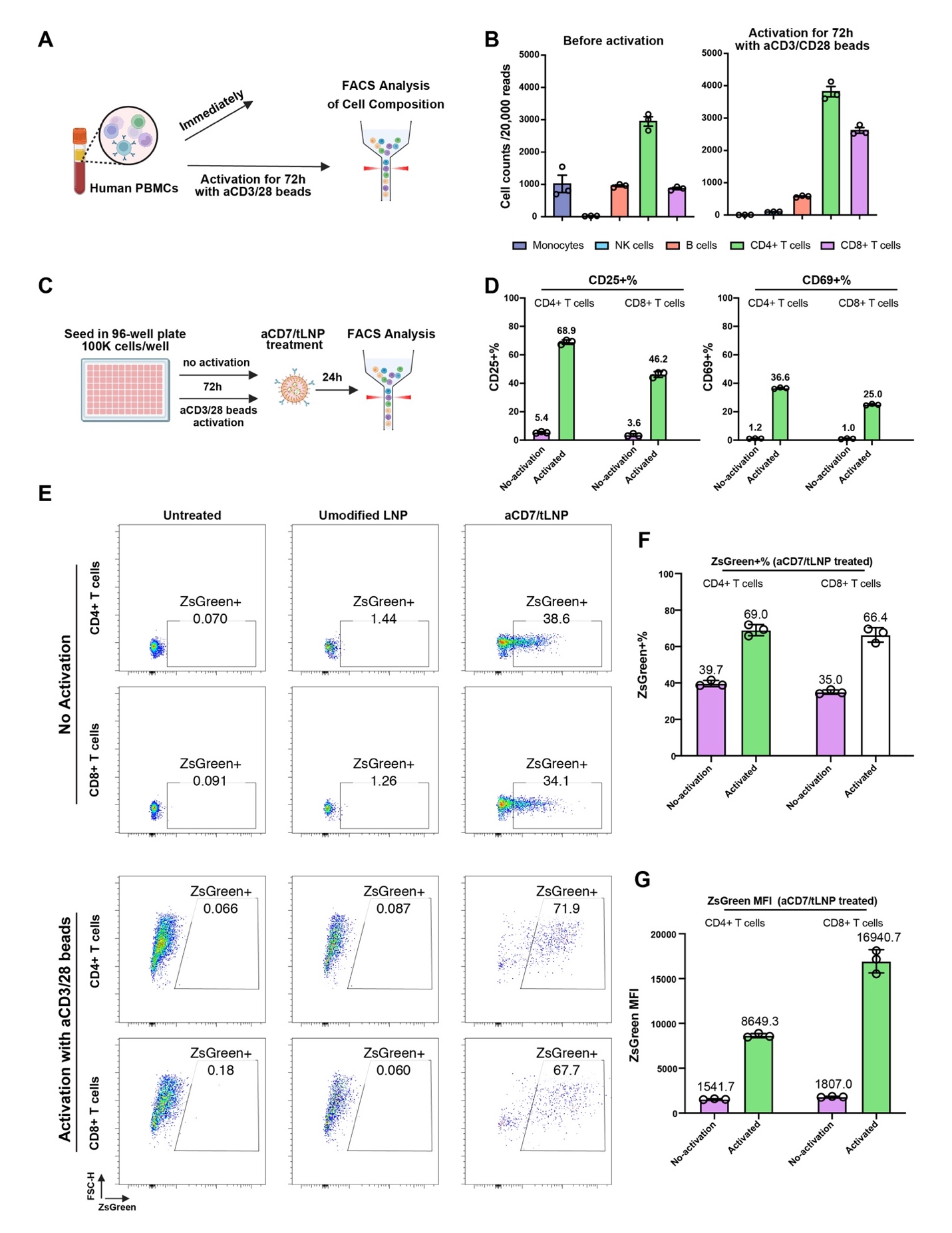

**Fig. S3. Activation of primary human T cells improves uptake and expression of mRNA delivered by CD7-targeted LNPs. (A)** Schematic for assessing immune cell composition in freshly isolated human PBMCs and after 72-hour activation with anti-CD3/CD28 beads. **(B)** Quantification of major immune cell subsets before activation (left) and after 72-hour activation (right). **(C)** Experimental design for evaluating aCD7/tLNP-ZsGreen delivery efficiency in resting or activated human T cells. **(D)** Expression of activation markers CD25 and CD69 on CD4⁺ and CD8⁺ T cells under non-activated and activated conditions. **(E)** Representative flow cytometry plots showing ZsGreen reporter expression in CD4⁺ and CD8⁺ T cells treated with unmodified LNP or aCD7/tLNP at 1 μg mRNA per 100,000 cells dosage, comparing non-activated (top) and activated (bottom) conditions. **(F)** Percentage of ZsGreen⁺ CD4⁺ and CD8⁺ T cells following aCD7/tLNP treatment under non-activated or activated conditions. **(G)** Mean fluorescence intensity of ZsGreen signal in CD4⁺ and CD8⁺ T cells treated with aCD7/tLNP. Bars represent mean ± SD, with individual donor values shown. *n* = 3 biological donors.

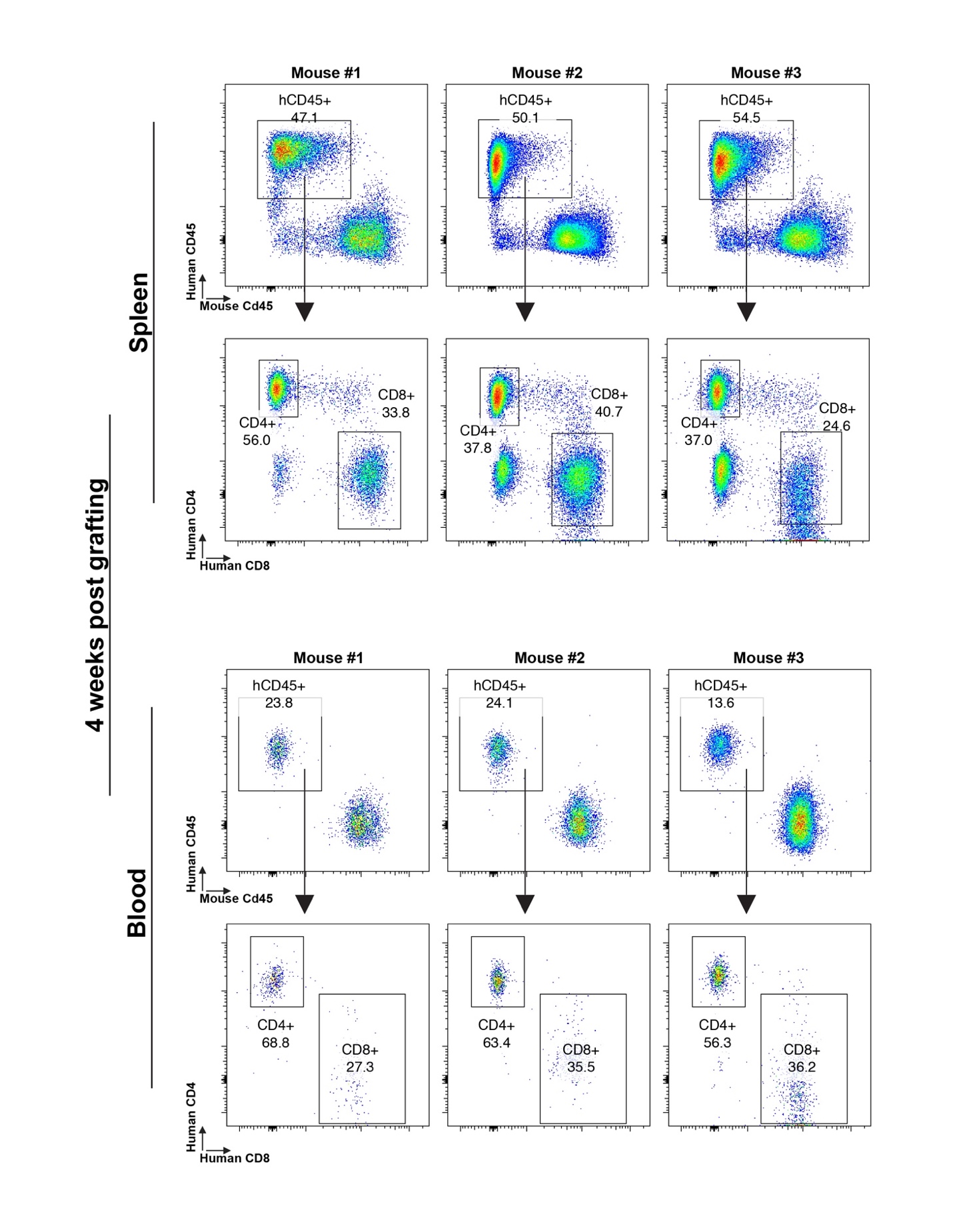

**Fig. S4. Verification of successful humanization of NSG mice 4 weeks post-grafting of human PBMCs by assessing human T-cell populations in peripheral blood via flow cytometry.**

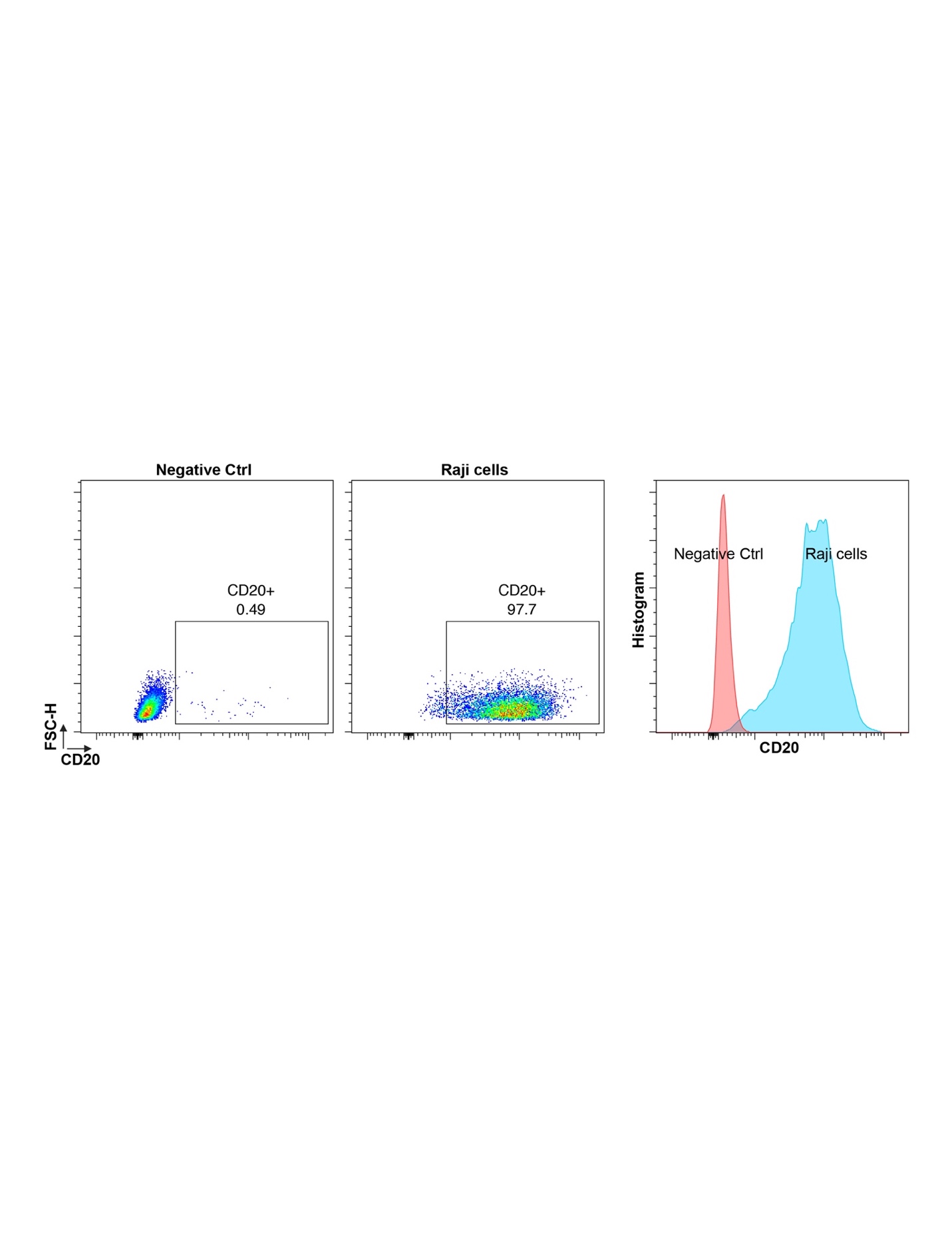

**Fig. S5. Verification of CD20 expression in the Raji cell line used as the target for aCD7/tLNP-mRNA generated CAR T cells.**

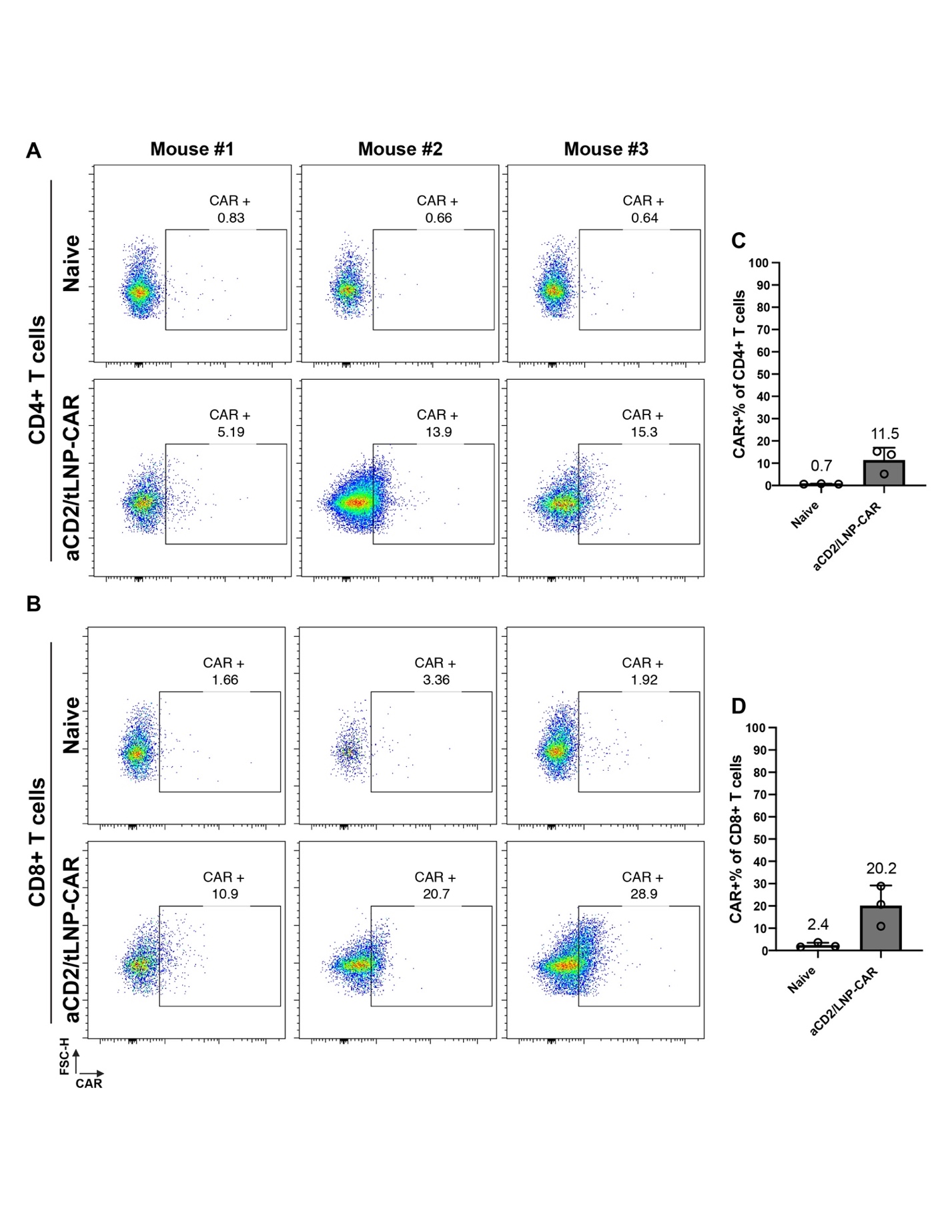

**Fig. S6. aCD2/tLNP-aCD20 CAR mRNA generates CAR T cells at a lower efficiency *in vivo*.** **(A-B)** CAR expression in CD4⁺ T (A) and CD8⁺ T (B) cells in the spleen 24-hours after treatment with aCD2/tLNP-CAR mRNA (2.5 μg per mouse), analyzed by flow cytometry. **(C-D)** Quantification of CAR expression in CD4⁺ T (C) and CD8⁺ T (D) cells. *n* = 3 mice, Data are shown as mean ± SD.
